# Generation of *Chlorella vulgaris* starch mutants and their biomass and lipid productivities under different culture media

**DOI:** 10.1101/2025.10.28.685034

**Authors:** Ana C. Ramos, Anna Hamilton, Isabel Molina, Patrick J. McGinn, Sharon Regan

## Abstract

**Background:** Microalgae are an important feedstock for the production of a wide variety of products, including biodiesel. Biodiesel, composed of fatty acid alkyl esters, is produced through the transesterification reaction of triacylglycerol (TAG). Microalgae store their energy reserves primarily as starch and TAGs. Therefore, several studies have focused on understanding the partitioning of carbon precursors between starch and TAG biosynthetic pathways. In this study, 5 starch mutants of *Chlorella vulgaris* were developed and cultured on different culture media.

**Results:** *Chlorella vulgaris* starch mutants were generated through UV-random mutagenesis. Five starch mutants were selected for this study: four low-starch producing mutants (st27, st29, st43 and st54) and one high-starch producing mutant (st80). The starch mutants were cultured on media with different organic carbon sources, and lipid and biomass productivity were measured. Mixotrophic growth on glucose resulted in the highest lipid productivity in all the mutants, including st80, without compromising growth, whereas photoautotrophic growth generally did not result in changes in lipid productivity of the starch mutants. The highest increase in lipid productivity was observed for st27, with a 3.8-fold higher lipid productivity than wildtype.

**Conclusions:** All starch mutants increased their lipid productivities when grown mixotrophically on glucose, suggesting the overflow hypothesis could explain the partitioning of carbon between starch and TAGs. Out of the mutants generated in this work, st27 resulted in the highest increases in lipid productivities, reaching an increase of 380% when grown mixotrophically on glucose, without compromising growth. The high-starch producing mutant st80 provides insight into a possibility to develop starch- and TAG-rich microalgal biomass.

## BACKGROUND

With an increasing need for sustainable processes, microalgal biomass has become an important feedstock as a wide variety of products can be obtained from these microorganisms (reviewed by Harun et al. 2010). As photosynthetic organisms, microalgae contain a variety of pigments and fatty acids that hold great interest for the cosmetic and nutraceutical industries, where products like lutein, astaxanthin and docosahexanoic acid are in high demand (Cordero et al. 2011, reviewed by Ethier et al. 2011, reviewed by Kim et al. 2016). Microalgal biomass has also become an important feedstock for the production of sustainable energy, where biodiesel (Přibyl et al. 2012), bioethanol (Jeon et al. 2013), biomethane (Mendez et al. 2013) and biohydrogen (Chader et al. 2011) can be obtained. Microalgal biodiesel is a mix of fatty acid methyl esters (FAMEs) produced after the extraction and transesterification of triacylglycerol (TAG), usually with methanol and an alkaline catalyst (reviewed by Halim et al. 2012). Thus, increasing TAG productivity in microalgae cultures has become a priority to increase biodiesel production from this feedstock (reviewed by Chisti 2007). It has been demonstrated that stress conditions, such as N-deficiency, result in an increase in TAG formation. However, N-deficiency leads to a significant decrease in biomass production, limiting the overall biodiesel productivity (Msanne et al. 2012, Allen et al. 2015, Benvenuti et al. 2015). To avoid the impact on productivities, several studies have focused on understanding the partitioning of carbon between energy reserves to increase metabolic flow towards TAG synthesis (Ramazanov and Ramazanov 2006, Work et al. 2010, Breuer et al. 2014).

Microalgae can grow on highly variable nutrient sources, including both oxidized and reduced nitrogen forms (reviewed by Cai et al. 2013, Ramanna et al. 2014). Different carbon sources can also be assimilated: microalgae can grow photoautotrophically (relying solely on photosynthesis) using CO_2_ as the inorganic carbon source, heterotrophically using diverse organic carbon sources such as glucose and acetate, and mixotrophically using both organic and inorganic carbon sources (Ren et al. 2014, Leite et al. 2015). Once the microalgae cells meet their metabolic demands, the remaining carbon can be stored into energy reserves to use in case environmental conditions become unfavorable (Msanne et al. 2012, Qian et al. 2014, Fan et al. 2015, Li et al. 2015). Just like higher plants, microalgae store energy mainly in two forms: starch and TAGs, with starch being more predominant (Fan et al. 2011, De Jaeger et al. 2014, Pancha et al. 2014). Therefore, carbon not immediately used for growth, energy or structure in a microalgal cell are stored into TAGs after starch metabolism is saturated (Fan et al. 2012, Fernandes et al. 2013). Several studies suggest that inhibition of starch formation results in a higher production of TAG in *Chlamydomonas reinhardtii* (Zabawinski et al. 2001, Work et al. 2010), *Scenedesmus obliquus* (Breuer et al. 2014, De Jaeger et al. 2014) and *Chlorella pyrenoidosa* (Ramazanov and Ramazanov 2006), and that without the option to synthesize starch the remaining alternative for the cell to store energy is through TAG. Thus, it is hypothesized that in the absence of a starch biosynthetic pathway the flow of carbon reserves is directed towards the synthesis of TAG, commonly referred to as the overflow hypothesis.

In the present study, *Chlorella vulgaris* starch mutants were developed through random mutagenesis with UV-C light to investigate if mutants with lower production of starch have increased TAG yields. Different culture media were used to compare growth and lipid production of the starch mutants in the presence of different nitrogen and carbon sources. This work provides further evidence on the overflow hypothesis in microalgae of the genus *Chlorella*, and provides evidence on the importance of the carbon source in the culture media for TAG production. To our knowledge, this is the first report of *Chlorella vulgaris* starch mutants.

## METHODOLOGY

### Microalgae strain and growth

The microalga *Chlorella vulgaris* (UTEX, B1803) was maintained in Bold’s Basal medium (BBM) in 250 mL Erlenmeyer flasks with a culture volume of 50 mL. The cultures were continuously shaken at 100 rpm, and incubated at 22 °C with continuous white light (20 µmol s^−1^ m^−2^).

### Random mutagenesis

*Chlorella vulgaris* mutants were developed after exposure to UV-C light (254 nm) as described previously (Vigeolas et al. 2012). Briefly, wild-type cultures were inoculated to an initial cell density of 5 x 10^5^ cells/mL in BBM at 100 rpm, 22 °C and continuous illumination. The culture was left to grow 2-3 days until it reached a cell density of 2-3 x 10^6^ cells/mL. Once, the culture reached the desired cell density, 25 mL of the algal suspension where transferred to an open sterile Petri plate with a sterile magnetic stir bar. The culture was then exposed to UV-C light (254 nm) for 13 min with the lamp (Handheld UVG-4, UVP) held 7.5 cm away from the culture to achieve a survival rate between 5-10%.

After exposure, the mutagenized cells were spread on Petri plates containing BBM-agar and incubated overnight at room temperature in the dark to avoid photorepair mechanisms. The plates with the mutagenized cells where then incubated at room temperature under continuous illumination until green colonies developed (approximately 7 days). The mutants were replica-plated on N-deficient BBM (BBM-N)-agar Petri plates and incubated for 10 days under constant illumination. Colonies on the replica plates were exposed to iodine vapor for 10 min to identify starch mutants as previously described (Work et al. 2010). To expose colonies to iodine vapor, solid iodine crystals were placed on the surface of an airtight chamber containing the uncovered replica plates with the mutants. Mutants that stained indigo/dark blue were considered starch producing, whereas mutants that stained brown/orange or did not change in color were considered low-starch mutants (starchless). Approximately 85 putative starch mutants were produced, and 5 were selected for further analysis because of their stable phenotype.

### Cultivation conditions and harvest

Starch mutants were maintained in both liquid and solid BBM under continuous illumination. Several culture media were tested: BBM, BBM supplemented with 15 g/L glucose (BBM+gluc), tris-acetate-phosphate (TAP) medium, and tris-phosphate (TP) medium. The mutants were grown on 10 mL of each of these media for a period of 7 days for acclimation in 50 mL Erlenmeyer flasks at 22°C under continuous illumination and 100 rpm. After the acclimation period, each culture was inoculated to an initial cell density of 10^5^ cell/mL in 250 mL Erlenmeyer flasks with 50 mL culture volumes, and incubated at 22 °C under continuous illumination and 100 rpm over a period of 8 days. To trigger TAG synthesis, N-deficiency was also compared: the cultures were grown for 5 days on BBM, harvested, and resuspended in 50 mL of BBM without nitrate (BBM-N) for the remaining 3 days of cultivation. Biological triplicates were studied for all the culture media tested.

### Confirmation of low-starch phenotype

Given that there is no selective pressure to maintain the mutation, the mutants’ phenotype was monitored periodically to confirm the phenotype was maintained. As a quick confirmation of the low-starch phenotype, 10 mL of each microalgal culture were harvested by centrifugation at 3000 x g for 20 min in a 15 mL conical tube. An iodine solution was prepared by adding 5 mM I_2_ and 5 mM KI in water. One mL of the iodine solution was added to the microalgal pellets and incubated at 75°C for 10 min in a water bath. To optimize starch staining, 1 mL of hot iodine solution was added and the tubes were vortexed. Starch phenotypes where determined by the intensity of the blue color after iodine staining, where high abundance of starch was visually determined by an indigo color.

To corroborate the starch-less phenotype at a cellular level, 150 µL of each culture at late exponential phase were harvested by centrifugation at 15,000 x g for 10 min in a 500 µL micro-centrifuge tube. The pellets were resuspended in 20 µL of the iodine solution and incubated at 90 °C for 10 min. Samples were left to cool down to room temperature, and visualized with bright-field microscopy under 1000X magnification (Axio Imager Z.1, Zeiss). *Chlamydomonas reinhardtii* starchless mutant BAFJ5 (Zabawinski et al. 2001) was used as a control to corroborate phenotypes through iodine staining.

### Analysis of starch content in microalgal biomass

To determine the amount of starch produced by the starch mutants relative to wildtype, a colorimetric semi-quantitative assay was employed. The assay was developed from a combination of protocols from previous studies (Xiao et al. 2006, Fernandes et al. 2012). Briefly, microalgal cultures were grown until late exponential phase; biomass was harvested by centrifugation at 15,000 x g for 10 min, and freeze-dried over 24 hrs. Dry biomass was then ground in a mortar and pestle using liquid-N until a fine powder was obtained. The ground biomass was then resuspended in 80% ethanol, and incubated at 75°C for 15 min to facilitate pigment extraction. Biomass was then precipitated by centrifugation at 17,600 x g for 10 min, and resuspended again in 80% ethanol. Pigment extraction was repeated several times, until ethanol extract was colorless. After removing ethanol, the ground biomass was resuspended in water to a concentration of 2 mg/mL. In a 96-well microplate, 20 µL of the resuspended biomass were added to a well containing 180 µL of iodine solution (5 mM I_2_ and 5 mM KI), mixed by pipetting up and down several times, and starch-iodine complex formation was recorded by optical density in a spectrophotometer (SpectraMax Paradigm, Molecular Devices) at 580 nm.

### Biomass production and growth analysis

The final cell density of the suspension was obtained by cell counting using an improved Neubauer hemacytometer (Bright-Line, SIGMA Aldrich) under a microscope with 400X magnification. Analytical duplicates were done for each biological replicate.

Biomass concentration was determined gravimetrically. 10 mL of the algal suspension were transferred to pre-weighted 15 mL polypropylene conical tubes. The cells were harvested by centrifugation at 3000 x *g* for 20 min, the supernatant was discarded, and the pellets were freeze-dried for 48 hr. Dry pellets were weighted to calculate biomass concentration. Growth of the mutant lines and wildtype is presented in terms of biomass productivity (BP, g_DW_ L^−1^ day^−1^), calculated as the product of their specific growth rate (µ, day^−1^) and biomass concentration (B, g_DW_ L^−1^) at the end of the cultivation period, following Equation 1.

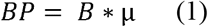

### Lipid profile

Fatty acid methyl esters (FAME) produced from the starch mutants and wild-type were obtained by the acid-catalyzed direct transesterification, using dilute sulphuric acid as the catalyst at high temperature. Briefly, dry biomass (approximately 2.5 mg) was transferred to a clean glass tube. Fifty microliters of a 2 mg/mL stock solution of tripentadecanoin (15:0 TAG) in 1:1 heptane:toluene was added as the internal standard to be used for quantification of the individual FAMEs produced. Each sample was dissolved in 0.5 mL of toluene and 1 mL of 5% sulphuric acid in methanol (v/v) was added. Then, 25 μL of 0.2% butylhydroxytoluene (BHT) was added to prevent unwanted oxidation reactions. The tubes were flushed with nitrogen gas, capped, and vortexed for 30 seconds. The tubes were heated at 80°C for 2 hours, with vortexing every 15-20 min for 15 sec. Samples were then cooled to room temperature and 1.5 mL of 0.9% sodium chloride solution was added, followed by 3 mL of hexanes to facilitate FAME extraction. The tubes were mixed by vortexing, then centrifuged at 300 x *g* for 2 min. The hexane layer was transferred to a clean tube. A second extraction using 3 mL of hexanes added to the lower layer was done, and the aqueous phase was discarded. Anhydrous sodium sulphate (Na_2_SO_4_) was added to the pooled hexanes, briskly agitated, and left overnight at room temperature. The tubes were centrifuged, the hexanes were transferred to a clean tube and the pooled extracts were dried under nitrogen. The dried extracts were dissolved in 1 mL hexanes and analyzed on a Thermo Scientific TRACE 1300 gas chromatograph coupled to a flame ionization detector (GC/FID) equipped with a TG-5MS capillary column (Thermo Scientific; 30 m length, 0.25 mm i.d., and 0.25 mm film thickness) using split injection (25:1 ratio, 310 °C). The oven temperature was programmed to increase from 120 °C to 280 °C heating at 3 °C per minute, with a final five minute-hold at 280 °C, and using a helium gas flow of 1.5 mL/min. Individual FAMEs were identified by comparison to the retention times of reference standards and confirmed by GC-MS. Peaks were quantified based on FID ion current.

Lipid productivity (LP, µg_FAME_ L^−1^ day^−1^) was calculated as the product of the total FAME content in biomass dry weight (DW) (L, µg_FAME_ g_DW_^−1^) and the BP (g_DW_ L^−1^ day^−1^), following Equation 2.

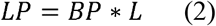

## RESULTS

Given that microalgae accumulate both starch and TAGs as energy reserves, the present study looked at the effect of inhibiting the starch biosynthetic pathway in *Chlorella vulgaris* had on lipid productivity. *Chlorella vulgaris* suspensions were exposed to UV-C light producing more than 7000 independent mutants, out of which 5 starch mutants (st) were selected to study after showing persistence of a phenotype over time: four low-starch mutants (st27, st29, st43 and st54) and one starch-overproducing mutant (st80). The phenotypes were determined by iodine staining (Figure 1), tentatively indicating that abundance of starch followed st80 > wild-type > st43 > st54 > st29 > st27. The phenotypes were corroborated through starch quantification in the mutant’s biomass, confirming that st27, st29, st43 and st54 cells have less starch than wild-type, and st80 produces higher amounts of starch (Figure 2). This analysis was conducted on biomass grown on BBM and BBM+glucose, both presenting the same pattern of starch content, indicating abundance of starch follows the order st80 > wild-type > st54 > st43 = st29 = st27. Observed phenotypes have been maintained for over 2 years.

**Figure 1.**
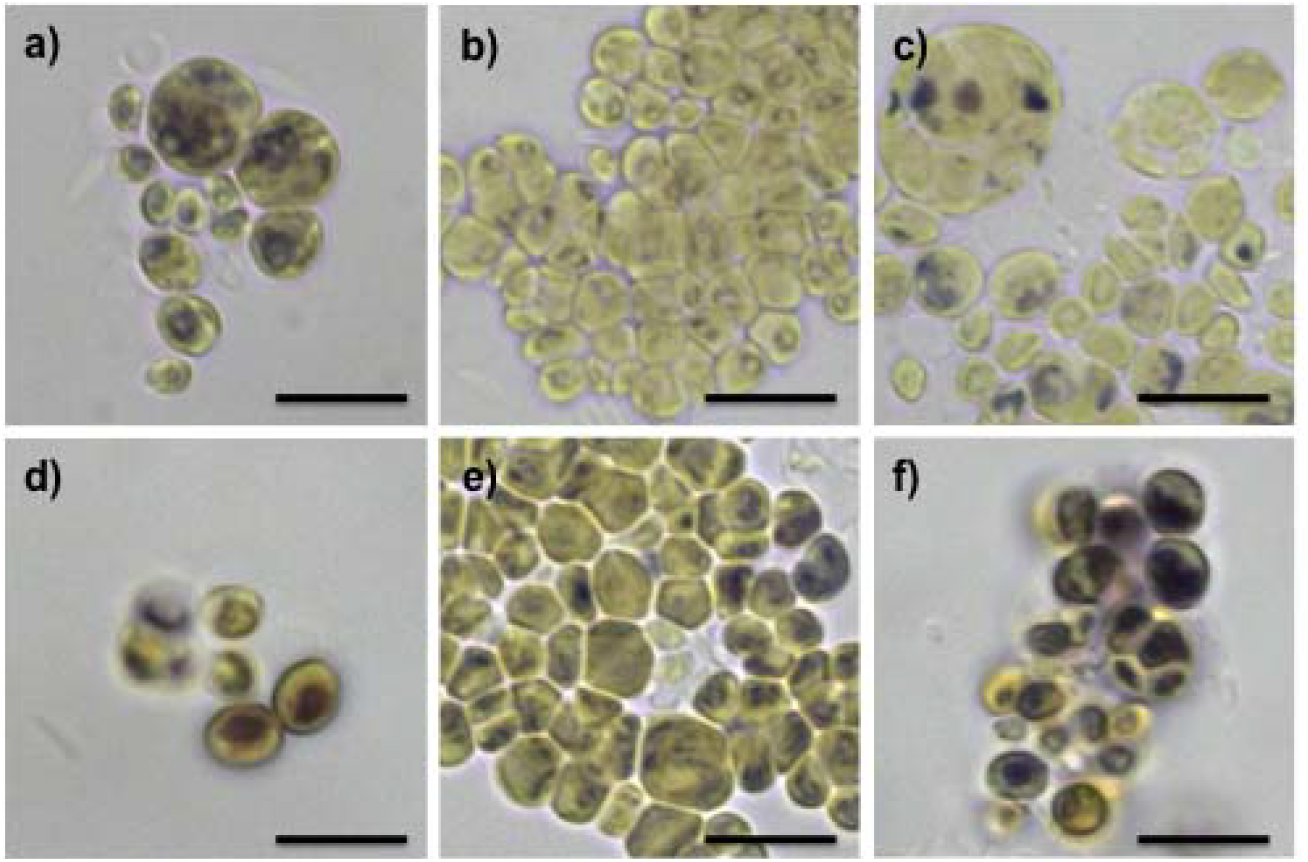
Iodine staining of *Chlorella vulgaris mutants* a) wildtype, b) st27, c) st29, d) st43, e) st54, and f) st80. Scale bar = 5 µm.

**Figure 2.**
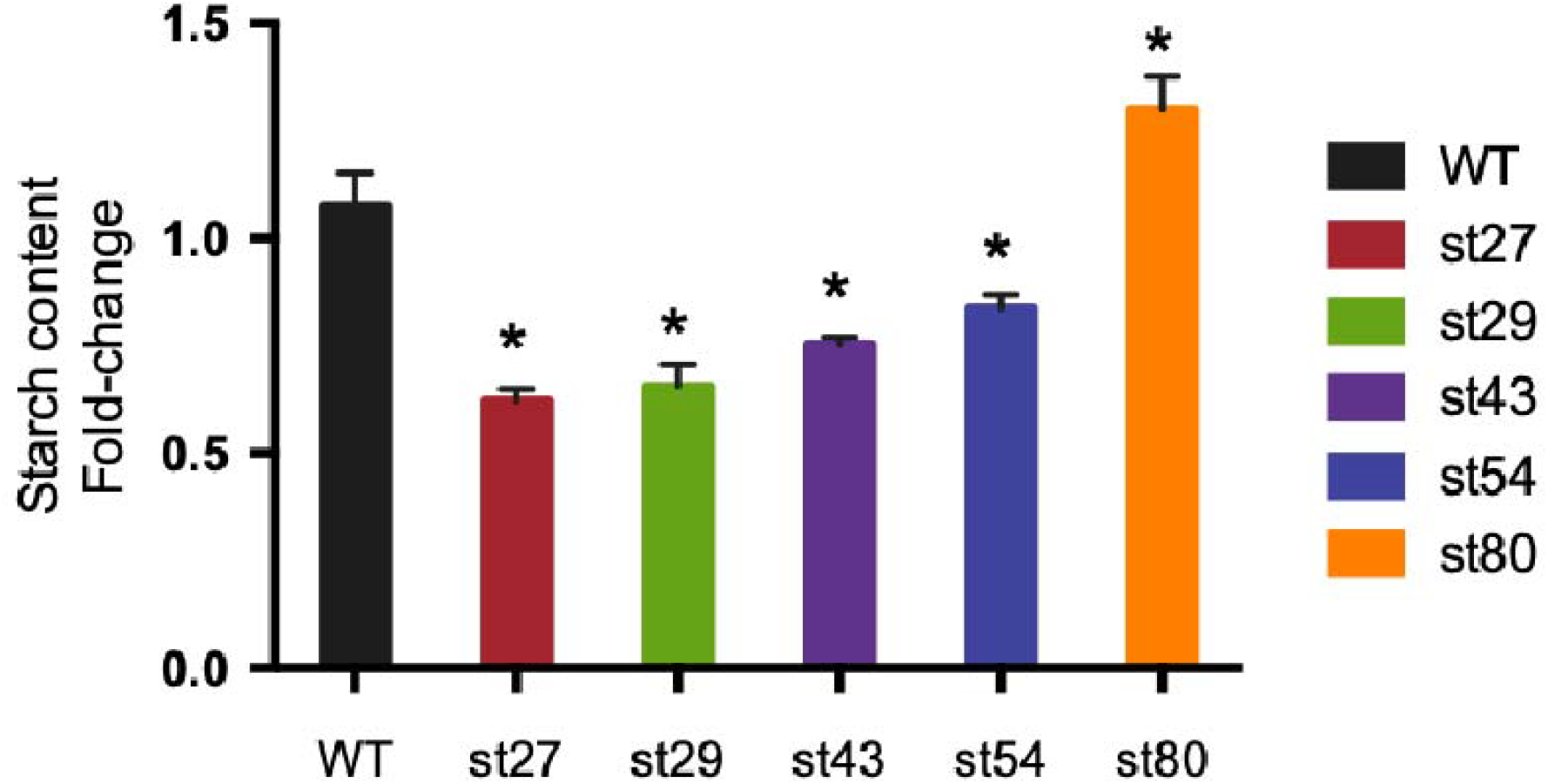
Colorimetric analysis of starch content in mutants relative to wildtype. Errors bar indicate standard deviation (n = 3). Starch content in all the mutants was significantly different than wildtype (p < 0.0096).

The five generated mutants presented differential starch accumulation, as revealed by their coloration patterns after iodine staining. The wildtype had indigo coloration typical of a starch-iodine complex formation at a localized point surrounding the pyrenoid, as well as randomly spread throughout the cell (Figure 1.a); in contrast, st27 produced less indigo coloration, mostly observed around the pyrenoid (Figure 1.b). Mutant st29 revealed a wide variation in cell sizes, where some cells do not present any indigo coloration (Figure 1.c). Interestingly, a brown stain resulted after exposing st43 to iodine, and the coloration appeared very defined and localized (Figure 1.d). The low-starch producing st54 did not show major differences in the coloration pattern, but the intensity of the indigo coloration appears weaker in st54 than in wildtype (Figure1.e), whereas st80 presented a very strong indigo coloration, indicative of higher starch content (Figure 1.f).

### Growth of starch mutants is mostly affected by glucose consumption

Given that biodiesel yield is directly proportional to both lipid and biomass production of the microalgal feedstock, biomass productivity and growth rates were quantified in this study (Figure 3 and Table 1).

**Table 1.**
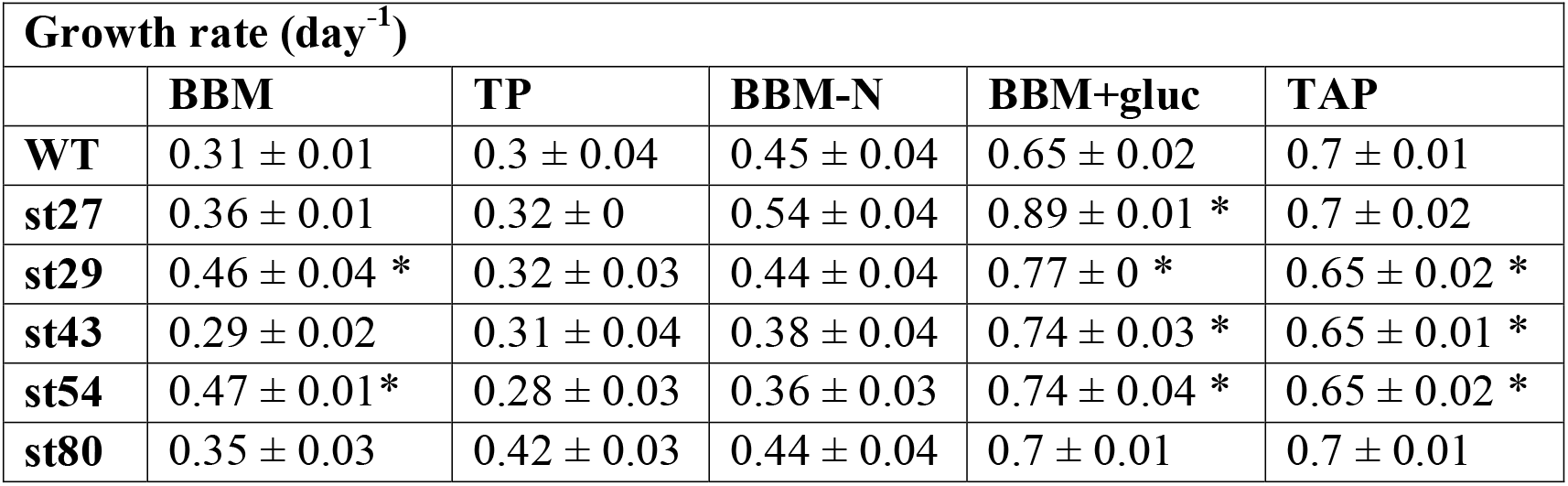
Calculated specific growth rates by the different culture media, expressed as average ± standard deviation (n = 3). Asterisk indicates statistical significance when compared to wild-type (p-value < 0.01321).

**Table 2.**
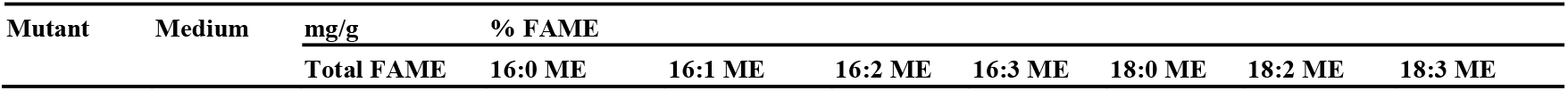
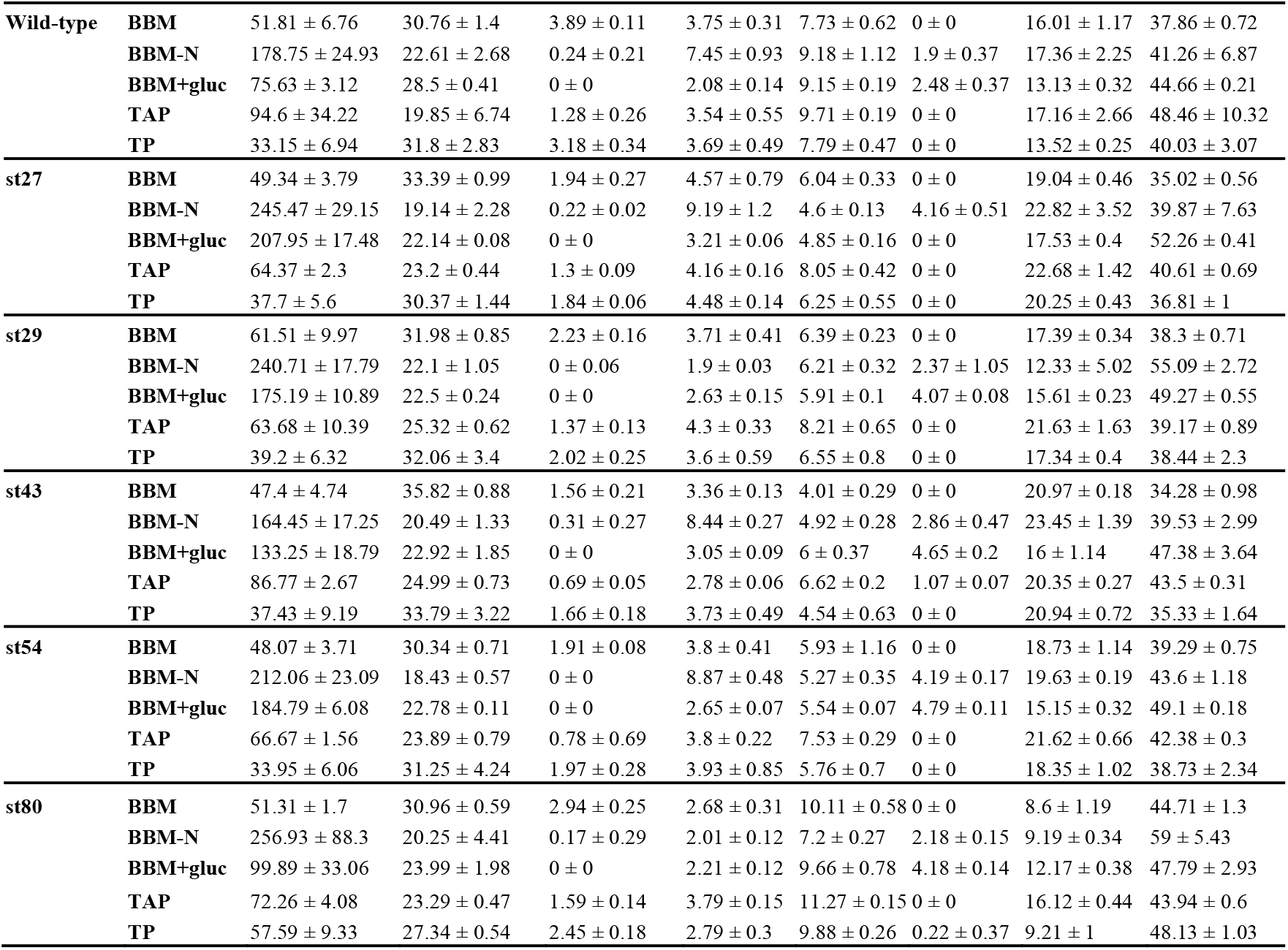
Lipid profile of *Chlorela vulgaris* starch mutants grown on different photoautotrophic (BBM, TP and BBM-N) and mixotrophic (BBM+glucose and TAP) culture media (n=3). FAME = fatty acid methyl ester.

**Figure 3.**
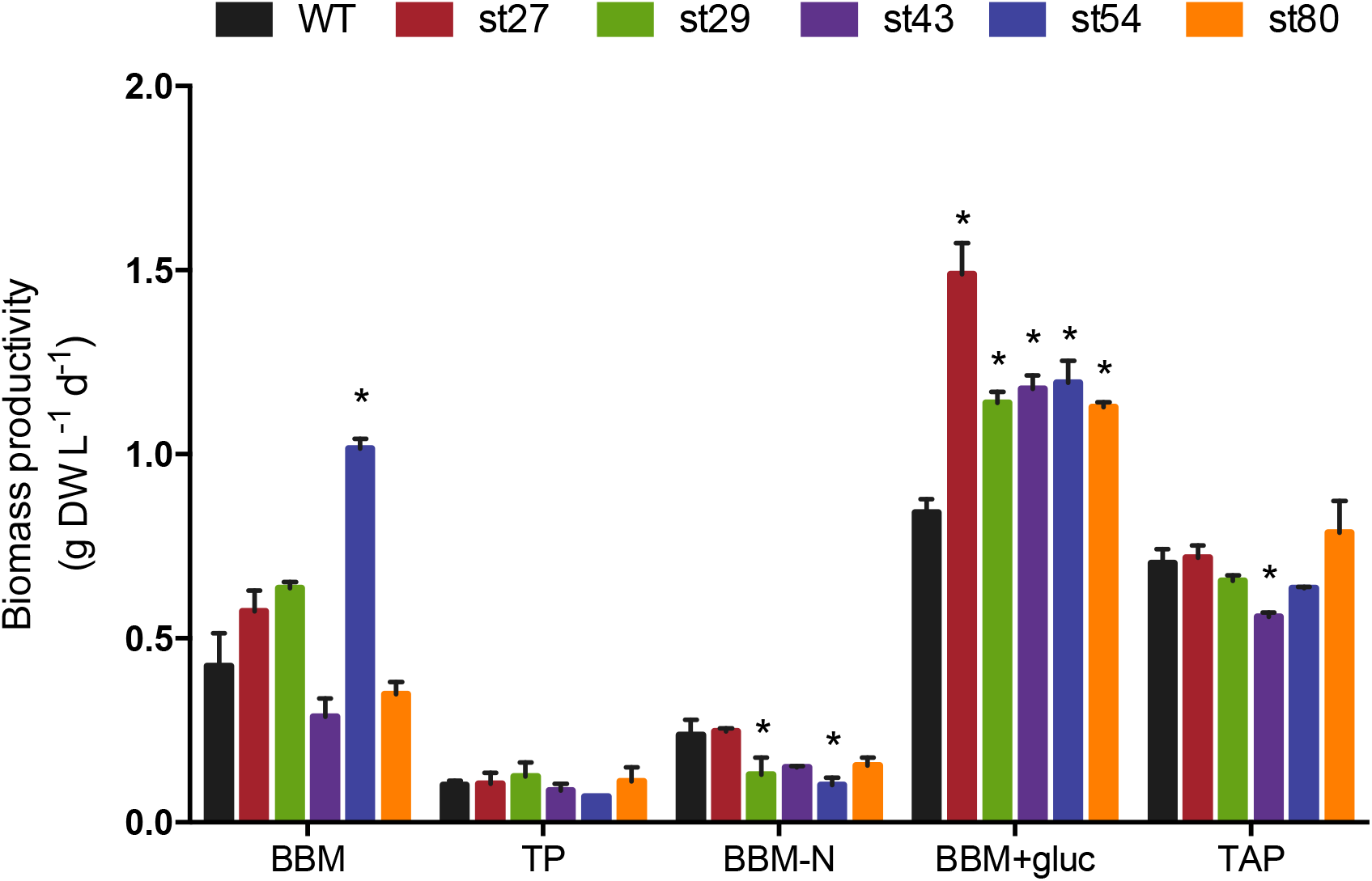
Biomass of *Chlorella vulgaris* starch mutants grown photoautotrophically (BBM, TP and BBM-N) and mixotrophically (BBM+glucose and TAP). Error bars indicate standard deviation of biological triplicates. Asterisks indicate significance compared to wild-type (p-value < 0.04438).

Photoautotrophic cultivation did not produce any changes in biomass productivity, with the exception of st54 grown on BBM, where biomass productivity increased by 140%. As expected, all the tested lines had lower biomass productivities during photoautotrophic cultivation under N-limitation (BBM-N), consistent with findings from other studies where growth is negatively affected by this stress (Msanne et al. 2012, Praveenkumar et al. 2012, Vonlanthen et al. 2015). However, st29 and st54 showed a significantly lower biomass productivity than wildtype when grown on BBM-N, but their growth rates were not affected (Table 1).

Of the culture media tested, the strongest effects on biomass productivity were observed during mixotrophic cultivation on glucose. Growth on BBM+glucose resulted in an increase in biomass productivity in all the lines (p-value < 0.02351), with the highest increase of 76% observed in st27 (p-value = 0.00012) (Figure 3). Growth rates of all the low-starch mutants increased in BBM+glucose, but st80 did not result in any changes in growth rate (Table 1).

Growth on TAP medium did not result in significant changes in biomass productivity of the mutants, with the exception of st43, which decreased biomass productivity by 20% (p-value = 0.00938)

### Lipid productivity is influenced by the culture medium

To determine whether the carbon source had an effect on carbon partitioning between starch and TAG, the starch mutants were grown photoautotrophically with atmospheric CO_2_, under N-deficiency, and mixotrophically with glucose and acetate. Lipid productivity was assessed in terms of FAME produced by day per volume of culture (Figure 4). When grown photoautotrophically, only st43 showed a decrease in lipid productivity by 33% on BBM (p-value < 0.01473), and a decrease by 29% on TP (p-value = 0.00031). Interestingly, st29 and st80 doubled their lipid production when grown on TP (p-value = 0.00021). As expected, growth on BBM-N caused an overall increase in lipid production in wildtype and the starch mutants; however, only st27 and st80 showed higher productivities than wildtype, resulting in 130% and 88% increases in lipid productivities, respectively (p-value < 0.00028).

**Figure 4.**
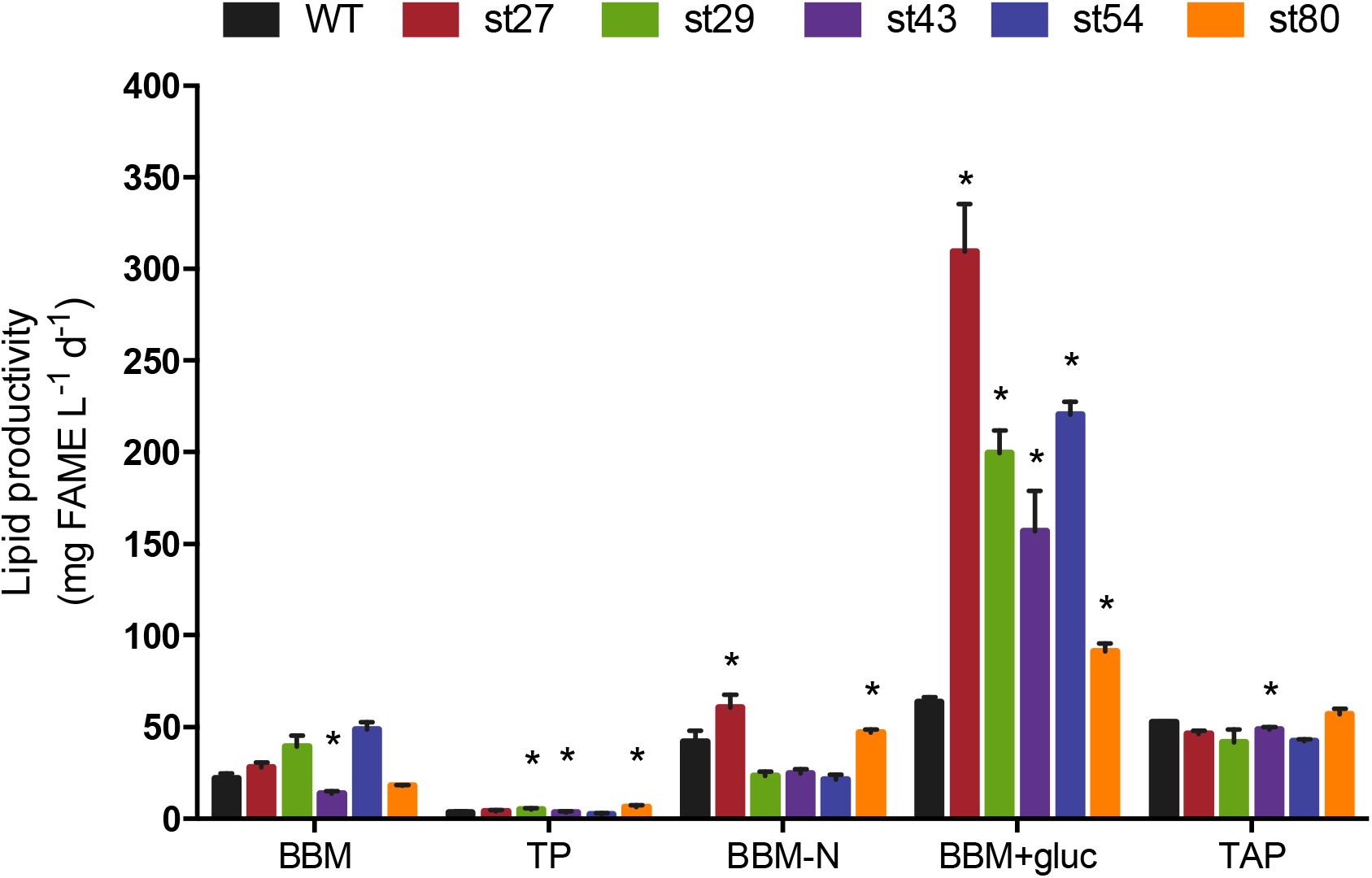
Lipid production of *Chlorella vulgaris* starch mutants grown on different culture medium. Error bars represent standard deviation of biological triplicates. Asterisks indicate significance compared to wild-type (p-value < 0.03433). FAME = fatty acid methyl ester.

Growth on TAP medium did not increase lipid yields in any of the starch mutants, including st80, and st43 even showed a slight decrease in lipid productivity (p-value = 0.03433). Growth on BBM+glucose, however, resulted in an increase in lipid productivity of all the low-starch mutants by at least 85% (p-value < 0.00442), with st27 reaching the highest lipid productivity 380% higher than wildtype (p-value = 0.00015) (Figure 4), and the lowest increase of 85% (p-value = 0.00442) in st43. The mutant st80 also showed an increase in lipid productivity during mixotrophic growth on BBM+glucose, which was unexpected as higher starch production would be expected to result in less available carbon for TAG synthesis (Figure 4).

## DISCUSSION

Starch mutants have been generated for *Chlamydomonas reinhardtii* (Zabawinski et al. 2001), *Scenedesmus obliquus* (De Jaeger et al. 2014), *Dunaliella tertiocolecta* (Sirikhachornkit et al. 2016), *Chlorella pyrenoidosa* (Ramazanov and Ramazanov 2006), *and Chlorella sorokiniana* (Vonlanthen et al. 2015), all resulting in variable efficiencies with regards to carbon partitioning towards TAG. To our knowledge, this is the first report of *Chlorella vulgaris* starch mutants, as well as the use of several culture media to investigate if the various organic carbon sources result in different degrees of lipid accumulation in the generated mutants. In this study, a random mutagenesis approach was selected over direct genetic engineering for several reasons. First, we were not able to replicate any of the stable genetic transformation protocols available for *Chlorella vulgaris* (Kim et al. 2002, Cha et al. 2012, Fan et al. 2015, Yang et al. 2015). Second, a random mutagenesis approach was chosen to generate a range of phenotypes within our starch mutants that were revealed by their varying responses to the culture media tested.

It has been suggested that inhibition of the starch biosynthetic pathway could lead to an increase in lipid production due to an overflow of resources into TAG synthesis, referred to as the overflow hypothesis hereinafter. Several starch-less strains make a strong case for this overflow hypothesis, including *Chlamydomonas reinhardtii* BAFJ5 (Zabawinski et al. 2001) and *Scenedesmus obliquus* slm1 (Breuer et al. 2014, De Jaeger et al. 2014). However, a study conducted by Siaut et al. (2011) challenged this hypothesis after demonstrating that lipid production of their starch-less mutants did not increase over the parent starch-producing strain (Siaut et al. 2011). The authors compared lipid production of different *Chlamydomonas starchless* (*sta*) mutants with inhibition of starch synthesis at different levels, including BAFJ5 (*sta6;* deficient in ADP-glucose pyrophosphorylase) and BAFJ6 (*sta7-1;* deficient in isoamylase) generated through random insertion of pARG7 (Libessart et al. 1995, Zabawinski et al. 2001, Posewitz 2004). Additionally, *sta7*-complemented strains show an increase in accumulation of both starch and lipids (Work et al. 2010), suggesting that inhibition of the starch synthesis is not pivotal to increase lipid accumulation. In contrast, *sta6*-complemented strains showed starch and lipid accumulation comparable to that of wildtype (it is unclear which reference strain was compared as wildtype here) (Li et al. 2010), suggesting that the point at which starch synthesis is impaired plays a role in the effect on TAG accumulation. In another study, three starch-less mutants of *Chlorella sorokiniana* did not show an increase in lipid production; such mutations affected genes encoding isoamylase and starch phosphorylase, two key enzymes involved in starch biosynthesis, but did not result in carbon partitioning into TAG biosynthesis (Vonlanthen et al. 2015). However, isoamylase and starch phosphorylase are involved in starch branching and do not play a role in partitioning of carbon into amylose nor amylopectin to form starch. Thus, the overflow hypothesis could still explain partitioning of carbon precursors between starch and TAG biosynthetic pathways.

In previous studies looking at carbon partitioning between starch and TAGs, increases in TAGs were explained by the availability of storage space in the chloroplasts (Fan et al. 2012, Li et al. 2015). This space-based theory proposes that, since both starch and TAGs are synthesized and partially stored in the chloroplast, the available storage space in this organelle is mostly occupied by starch (being the dominating carbon sink), leaving little space for TAG lipid bodies, thus decreasing TAG production in the cell (Fan et al. 2012, Li et al. 2015). This was supported by a study using *Chlorella sorokiniana*, where N-deficiency led to the reallocation of carbon from starch into TAGs (Li et al. 2015). The authors suggest that starch degradation is necessary for partial TAG synthesis to occur in this microalga during N-deficiency (Li et al. 2015), which is also observed in *Chlorella zofingiensis* (Zhu et al. 2014), likely to release space for TAG accumulation. This is not, however, what was found in *Chlorella sorokiniana* starch-less mutants ST86, ST3 and ST12 (Vonlanthen et al. 2015). These mutants, defective in isoamylase or starch phosphorylase, resulted in a reduction of starch between 60% and 94%, but did not cause any increases in FAME content; in fact, ST12 (defective in starch phosphorylase) had a decrease of 20% in FAME content during cultivation in N-deficiency (Vonlanthen et al. 2015). These *Chlorella sorokiniana* starch-less mutants, ST86, ST3 and ST12, produced up to 96% less starch than wildtype, suggesting that physical space in the chloroplast was available for TAG accumulation; yet, TAG content did not increase in these mutants, suggesting that the availability of physical space in the chloroplast is not a requirement to trigger TAG synthesis, but releasing carbon molecules from starch could be.

Results obtained in the present study support the overflow hypothesis, where increases in TAG content of starch-less mutants are explained due to an increase in carbon precursors available for TAG biosynthesis. This study included mixotrophic growth using glucose as the organic carbon source because glucose can be readily stored into starch reserves when metabolic needs are satisfied in the cell. Therefore, in the absence of an efficient starch biosynthetic pathway, extracellular glucose in lines st27, st29, st43 and st54 must be metabolized elsewhere. The fact that these low-starch lines increased lipid productivity and growth rates when grown on BBM+glucose suggests that carbon precursors normally allocated for starch synthesis become available for TAG synthesis and growth. The high-starch mutant st80 also increased lipid productivity when grown on BBM+glucose, without compromising biomass productivity or growth rate, suggesting that the high amounts of starch present in this line are not competing for space with TAG lipid bodies, in contrast to what the space-based hypothesis proposes. The increased levels of starch in st80, as well as the increased lipid productivity of this mutant on BBM+glucose, opens the possibility to generate microalgae strains capable of accumulating large amounts of both starch and TAGs, which would make this feedstock ideal for the recovery of various products (i.e. biorefinery applications).

The mutant lines generated in this study had different responses in the culture media tested, suggesting that the carbon source available in the culture medium has an effect on TAG production. In *Chlamydomonas reinhardtii* it was observed that starch is synthesized primarily using carbon derived from CO_2_ consumption, whereas consumed acetate is mostly incorporated into fatty acid synthesis (Juergens et al. 2016). Similarly, heterotrophic growth of *Chlorella vulgaris* supplemented with 1% acetate or glucose demonstrated that using acetate as the sole carbon source favored production of lipids and proteins over carbohydrates, whereas heterotrophic growth on glucose as the sole carbon source favored the production of carbohydrates, and provided the highest biomass production (Liang et al. 2009). Most studies on the available starch-less mutants in microalgae only explore one organic carbon source, typically acetate, so it is possible that different effects could be observed when testing various culture media.

In the present study, photoautotrophic cultivation did not result in a significant increase in lipid productivity in most of the starch mutants, suggesting that atmospheric CO_2_ consumed by these lines was not enough to satisfy metabolic demands and have a surplus to store as TAG. Similar findings were observed in *Dunaliella tertiolecta* starch mutants, where changes in TAG content could not be detected in the mutants when grown photoautotrophically under N-replete condition (Sirikhachornkit et al. 2016). Interestingly, in our study, growth on TP doubled lipid productivities of st29 and st80, without compromising growth, suggesting these lines may have higher CO_2_ fixation rates.

Mixotrophic growth on acetate (TAP medium) did not show a significant difference in neither biomass nor lipid productivities of the mutants (compared to wildtype), with the exception of st43. Multiple studies have come to the conclusion that overexpressing enzymes in the fatty acid biosynthetic pathway does not result in an increase in TAG production (Dunahay et al. 1996, Vigeolas et al. 2007, Fan et al. 2012, Ramanan et al. 2013, Li et al. 2015), even though TAG synthesis relies on the *de novo* fatty acid synthesis (Fan et al. 2011, Allen et al. 2015, Wang et al. 2015). Therefore, it is not surprising to see that st27, st29, st54 and st80 did not show significant changes in lipid productivity when cultivated on TAP medium, since acetate is mostly incorporated into fatty acid synthesis, which does not increase TAG production. An interesting observation is that st43 had a decrease in both biomass and lipid productivities when grown on TAP medium (20% and 30% respectively), and a decrease in lipid productivity in both N-replete photoautotrophic media (30%), but no changes when grown on BBM-N, suggesting that the TAG biosynthetic pathway is not affected, but this mutant may be preferentially allocating carbon in different pathways. Additionally, st43 showed the least increase in lipid productivity when grown on BBM+glucose compared to the other low-starch mutants. After iodine staining, st43 presented a brown coloration rather than the expected indigo coloration characteristic of an iodine-starch complex formation, suggesting a more prominent formation of other polysaccharides, most likely glycogen (Enjalbert et al. 2000, Vonlanthen et al. 2015). This tendency of st43 to accumulate other polysaccharides over starch could explain why this mutant produced more lipids than wildtype and st80 on BBM+glucose, but not as high as the rest of the low-starch mutants, as well as the decrease in lipid productivities when grown on TP and TAP. Therefore, it is possible there is less TAG synthesis in st43 because it is directing carbon towards synthesis of non-starch polysaccharides like glycogen. Similar findings were observed in *Chlorella sorokiniana* starchless mutants (Vonlanthen et al. 2015), but further research is needed to determine if st43 is a glycogen-accumulating mutant.

Future studies characterizing the mutations in these lines would assist in understanding how to efficiently trigger TAG accumulation in *Chlorella vulgaris*, without compromising other pathways. Identifying where the mutations occur to cause these starch phenotypes could provide valuable information in the generation of microalgal crops for additional products. For example, understanding where the mutation occurs in st43, as well as the polysaccharides it preferentially produces, could allow for the generation of a microalgal crop for the production of fermentable sugars. Furthermore, determining whether st80 has different CO_2_ fixation rates could provide valuable information for the conversion of the abundant greenhouse gas into high levels of starch and TAGs in *Chlorella*’s biomass.

## CONCLUSION

Five starch mutants were generated in this study: four low-starch producing mutants (st27, st29, st43 and st54) and one high-starch producing mutant (st80). All starch mutants increased their lipid productivities when grown mixotrophically on glucose, suggesting the overflow hypothesis could explain the partitioning of carbon between starch and TAGs. Out of the mutants generated in this work, st27 resulted in the highest increases in lipid productivities, reaching an increase of 380% when grown mixotrophically on glucose, without compromising growth. The high-starch producing mutant st80 may provide insight into developing starch- and TAG-rich microalgal biomass.

## DECLARATIONS

### Authors’ contribution

ACR designed experiments, maintained the microalgal cultures, generated the mutants, collected data for growth and starch content, interpreted the data, and wrote the manuscript. AH performed lipid extraction and FAME quantification. IM assisted in lipid analyses and edited the manuscript. PJM and SR assisted in experimental design, interpretation of the data, and edited the manuscript.

